# Adiposity QTL *Adip20* decomposes into at least four loci when dissected using congenic strains

**DOI:** 10.1101/171777

**Authors:** Cailu Lin, Brad D. Fesi, Michael Marquis, Natalia P. Bosak, Anna Lysenko, Mohammed Amin Koshnevisan, Fujiko F. Duke, Maria L. Theodorides, Theodore M. Nelson, Amanda H. McDaniel, Mauricio Avigdor, Charles J. Arayata, Lauren Shaw, Alexander A. Bachmanov, Danielle R. Reed

## Abstract

An average mouse in midlife weighs between 25 and 30 g, with about a gram of tissue in the largest adipose depot (gonadal), and the weight of this depot differs between inbred strains. Specifically, C57BL/6ByJ mice have heavier gonadal depots on average than do 129P3/J mice. To understand the genetic contributions to this trait, we mapped several quantitative trait loci (QTLs) for gonadal depot weight in an F_2_ intercross population. Our goal here was to fine-map one of these QTLs, *Adip20* (formerly *Adip5*), on mouse chromosome 9. To that end, we analyzed the weight of the gonadal adipose depot from newly created congenic strains. Results from the sequential comparison method indicated at least four rather than one QTL; two of the QTLs were less than 0.5 Mb apart, with opposing directions of allelic effect. Different types of evidence (missense and regulatory genetic variation, human adiposity/body mass index orthologues, and differential gene expression) implicated numerous candidate genes from the four QTL regions. These results highlight the value of mouse congenic strains and the value of this sequential method to dissect challenging genetic architecture.

---

An average mouse at six months of age weighs about 25 to 30 g. Some of this weight is due to adipose tissue located in several depots, the largest of which is the gonadal (Figure 1). In rodents, depots can differ in weight up to 20-fold, depending on the particular strain, and while adipose depot weight is highly heritable [1, 2], specific genetic determinants for each depot reflect its specialized functions [3]. Investigators have interbred specific strains and looked for chromosomal regions that affect the weight of individual adipose depots, and this procedure has identified hundreds of influential genomic regions, or quantitative trait loci (QTLs), which curators catalog in the Mouse Genome Database [4]. While it has been difficult to identify the underlying genetic variants for many complex traits, including adiposity [5], it remains an important scientific goal because this knowledge may better explain genetic risks and suggest new biological pathways and therapeutic targets [6].

**Figure 1.**
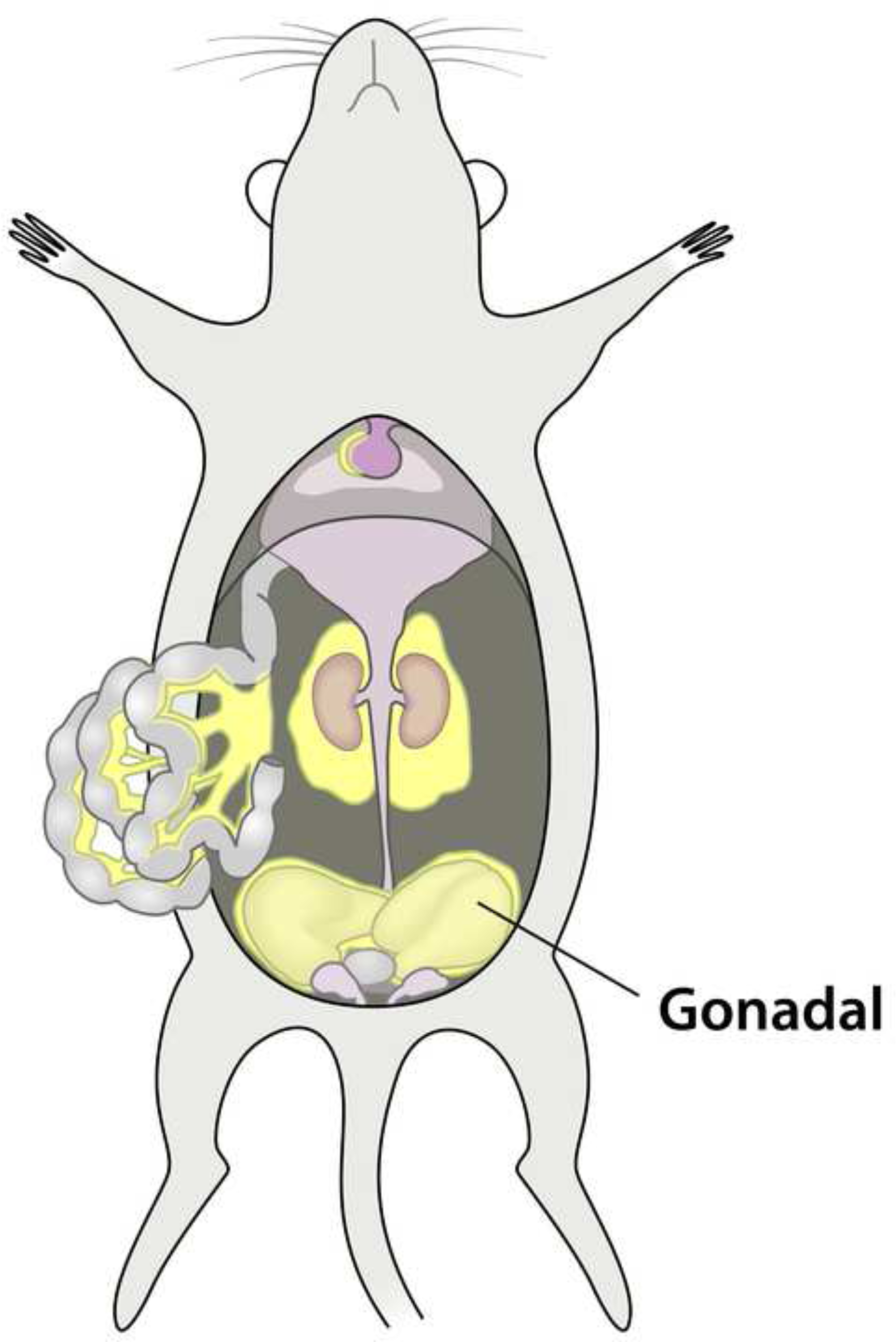
Anatomical location of the gonadal adipose depot in a male mouse.

Our specific goal here was to map QTLs for gonadal adipose depot weight in mice and to identify the underlying genetic variants using a pair of contrasting inbred strains. We began QTL mapping by intercrossing the C57BL/6ByJ (B6) strain with the 129P3/J (129) strain [7–9] and found a QTL on chromosome 9, which we originally called *Adip5* (at ~50 Mb) but was renamed *Adip20*. Chromosomal mapping using intercross populations did not provide sufficient resolution to narrow the genomic interval to specific candidate genes, so our next step was to refine the location of this QTL by creating and studying mouse congenic strains. We assayed or imputed genotypes for a common set of markers on mouse chromosome 9 and analyzed the genotype-phenotype associations using several computational approaches, including a sequential method of strain comparison [10]. Maternal effects and imprinting affect body composition, including adipose depot weight, so we incorporated several experimental design features to control these influences. After the mapping studies, we used several methods to identify candidate genes.

## Method

*Animal husbandry*. We bred all mice in the animal vivarium at the Monell Chemical Senses Center, located in Philadelphia, Pennsylvania (USA), except for the inbred B6 and 129 mice used for the first stages of breeding, purchased from the Jackson Laboratory. During this time, husbandry practices were stable, owing in part to having the same vivarium personnel and in part to using a consistent type of bedding (Aspen Shavings, Northeastern Products Corp, Warrensburg, NY). We fed mice Rodent Diet 8604 (Harlan Teklad, Madison, WI), and they lived in a 12:12 light cycle, with lights off at 6 pm, barring unusual circumstances (e.g., weather-related power outages). The Monell Institutional Animal Care and Use Committee approved these procedures.

We started branching congenic mice by backcrossing N_6_ (F_1_ x B6) males (from heterozygous consomic B6.129-Chr9 mice [11] with a partial donor region from chromosome 9) with B6 females to produce mice with 129-derived donor chromosomal regions of various lengths. We named the strains with codes that reflect their lineage; for instance, all strains with the prefix 1 (e.g., 1.1) descended from the same male. The breeding goal was to identify male breeders with a donor region that overlapped the QTL location discovered in the original F_2_ intercross.

We bred congenic mice that had either zero or one copy of the 129-derived donor region by mating inbred B6 female mice with heterozygous congenic males. This approach reduced maternal effects (all mothers were the same genotype) and reduced imprinting effects (only fathers contributed the donor region). Each congenic mouse was potentially genetically unique (because the paternal donor region could shorten due to meiotic recombination). Therefore, we genotyped each congenic mouse to ensure we could define the donor region breakpoints. While most mice had full-length donor regions, some had shorter regions, which we refer to as “partial.”

### Genotyping

We evaluated simple sequence-length polymorphism markers (e.g., *D9Mit2*) by polyacrylamide gel electrophoresis after polymerase chain reaction amplification by locus-specific primers [12] in our laboratories. We assayed single nucleotide polymorphisms (SNPs) at three locations: at the Genotyping and RNA Analysis Core at the Monell Chemical Senses Center; at the Center for Inherited Disease Research (see Electronic Resources) as part of an NIH-funded genotyping supplement; or by a commercial vendor (LGC, Beverly, MA; formerly KBiosciences) as a fee-for-service. When assaying variants in the Monell genotyping core, we used primers and allele-specific dye-labeled probes (Life Technologies, Carlsbad, CA). Irrespective of genotype location, controls (blank samples, and genomic DNA from inbred progenitors and their F_1_ hybrids) were included in all genotyping assays, and we retested unlikely genotypes as needed. We list all markers tested and their base pair positions in **S1 Table**. We imputed missing data by tracing the parental origin of the marker alleles and assuming that no double recombination occurred between markers 26 Mb or less apart. We selected this 26 Mb criterion as the smallest distance between two observed recombinant events in the most densely genotyped populations we have studied [11]. We also genotyped 12 representative congenic mice to evaluate residual genetic background using the Mouse Universal Genotyping Array (www.neogen.com) with 7,851 SNP markers built on the Illumina platform. We tallied the numbers of heterozygous markers in the genetic background (excluding the targeted donor regions) and expressed them as a fraction of the total valid markers for an estimate of residual heterozygosity.

### Phenotyping measures

After euthanasia by carbon dioxide asphyxiation, we weighed the body of each mouse to the nearest 0.1 g. During necropsy, we removed the gonadal adipose depot using anatomic landmarks [13, 14] and weighed the wet tissue to the nearest 0.1 g.

### Gonadal adipose depot gene expression

We conducted two microarray experiments to quantify gonadal depot adipose tissue gene expression between congenic mice with and without the donor region (Experiments 1 and 2). In Experiment 1, we selected tissue from male mice from congenic strain 4. We chose this strain because the donor region captured the *Adip20* peak we previously identified [15] (42.6 to 58.3 Mb). Half of the mice were heterozygous for the donor region (129/B6; N=7), and half were littermates homozygous for the donor region (B6/B6; N=7). Experiment 2 replicated Experiment 1 except there were N=6 mice in each genotype group, for a total of 12 mice. We isolated RNA from each gonadal adipose depot and measured genome-wide exon-by-exon expression using the Affymetrix Mouse Gene 1.0 ST Array. Members of the University of Pennsylvania Microarray Core facility prepared the RNA, performed the chip hybridization, and did preliminary quality control steps following manufacturer directions.

### Data analysis

We took three steps to prepare for the statistical analysis of the gonadal adipose depot weights. First, we eliminated data from mice that (a) appeared to be sick (e.g., hunched posture) or (b) had tumors or other obvious gross morphology found during necropsy (**S2 Table**). Second, we determined whether the data were normally distributed using the *fitdistrplus* R package [16], log-transformed the data (**S1 Figure**) if needed, and confirmed this transformation was effective using the Kolmogorov-Smirnov test (**S3 Table**). Third, we evaluated the importance of body weight and age as covariates using Pearson correlation coefficients. We observed that body weight was highly correlated with adipose depot weight but that age, owing to the tight range (measured in days), was not (**S2 Figure**). Therefore, we used body weight but not age as a covariate for the subsequent analyses:

#### 1. Marker association analysis

We treated all congenic mice as one mapping population and conducted a general linear model analysis of the log-transformed gonadal adipose depot weight for each marker, using marker genotype and strain as fixed factors and body weight as a covariate, and using a type 1 (sequential) sum of squares. For the presentation of the data, we (a) computed marker association test statistics and converted the associated p-values to the negative logarithm with base 10, (b) calculated the genotype mean, and (c) calculated the effect sizes using Cohen's *D* [17] for the peak linked marker in the mapping population. We defined a 'significant' statistical threshold as an α level of 0.05 after Bonferroni correction for the number of markers (N=148; -log_10_(α/*N*)= 3.47) and a suggestive threshold as an α level of 0.63 (– log_10_(α/*N*) = 2.37) [18].

#### 2. Common segment method

The common segment method is based on the assumption that a chromosomal region of interest harbors one QTL. Therefore, in our case, if the QTL is within a congenic donor region, there is a difference in gonadal adipose depot weight between host and donor mice, whereas congenic strains that do not share that QTL-containing donor region have no phenotype. Using this method, we conducted two analyses, one narrow and one broad. For the narrow analysis, we considered only a subset of congenic mice, those mice with full-length donor regions that were also from large genotype groups (N mice ≥ 12/strain/genotype; 12 strains). For the broad analysis, we used nearly all congenic mice as described below. For the narrow and broad analyses, we grouped mice within each congenic strain by whether they had a host or donor region, and compared these two groups using a general linear model with the presence of the donor region and strain as fixed factors and body weight as a covariate. We conducted the post hoc Fisher’s least significance difference to compare the phenotype between genotype groups for each congenic strain, and p<0.05 was used as a significance level. Those strains with a significant difference between genotype groups we classified as *Adip20* positive (+), and those that were not different we classified as *Adip20* negative (-).

#### 3. Sequential method

The sequential method can detect single and multiple QTLs [19] using an established procedure to compare congenic strains in a predetermined order. To establish the strain comparison order, we constructed a minimum spanning tree (MST) based on the donor region size and location with a custom R-script following the previously published pseudocode [19] supplemented with the R software package *optrees* [20] and *igraph.* We compared heterozygous congenic mice (129/B6) to the group of all homozygous littermates (B6/B6). To make comparisons between mice of different genotypes, we used a general linear model followed by least-squares difference post hoc analysis with a statistical threshold of p < 0.05.

### DNA sequence variants and human homology

Using an online database [21], we identified all genes residing within the genomic coordinates of the QTLs defined in the sequential congenic analysis. We next used another online database [22] containing genomic variants among inbred mouse strains from a large-scale genome sequencing project [23, 24] to compare the genomic sequences within the defined QTL regions for strains (129P2/OlaHsd and C57BL/6J) closely related to parental inbred strains used in our study. We formatted these variants using an online tool [25], and we identified regulatory variants as well as coding variants (missense and stop codon gain/loss) with the potential to cause functional changes. We determined which coding sequence variants were likely to be functional using computer algorithms in the program Sorting Intolerant From Tolerant [26]. Eight SNPs were excluded from this analysis because we could find no unique identifier for the variant in the dbSNP database (Build 138) [27]. We also identified human genes and their variants associated with obesity that are located in the regions of conserved synteny with the mouse QTL regions by searching an online catalog of human genome-wide association results [28] with key words “adipose” and “body mass index”.

### Analysis of microarray data

We performed gene expression analysis using Partek^®^ Genomics Suite^®^ (version 6.6, build 6.16.0419) by importing files of raw intensities for each probe set, performing the post-import quality control analyses, and comparing genotype groups (host versus donor region) for differential gene expression [29–31]. We report the differential expression as the log2-fold change (corresponding to a 1.5-fold change) with an associated p-value (p<0.05) corrected for false discovery using the number of genes examined [32]. We selected this relaxed fold-change cutoff to capture the anticipated small differences in gene expression that often arise from allelic variation [33]. We present data from the relevant genes using a customized volcano plot [34] annotating the resulting graph using the *ggrepel* [35] and *ggplot2* [36] R packages.

We compared the results between Experiment 1 and Experiment 2 and report only those genes that consistently differed by genotype. We included genes within the entire congenic donor region (not just the smaller QTL regions as defined below), reasoning that gene regulatory elements can be located millions of base pairs from their protein-coding regions [37]. We also examined the genome-wide pattern of differential expression beyond the donor fragment region of the congenic strain 4. We took this additional step to confirm our expectation that more genes would differ inside than outside the congenic region. To that end, we identified the top 200 differentially expressed genes (those with the lowest p-values) and used their chromosomal locations to create an ideogram [38].

For all analyses listed above, we computed the statistical tests with R (version 3.2.0) and R-studio (version 0.99.489) and graphed the results using GraphPad Prism 6 (version 6.05; GraphPad Software, La Jolla, CA). All data are available for download on Github (https://github.com/DanielleReed/Adip20) and the Center for Open Science (osf.io/yeqjf). All mentions of mouse genomic coordinates refer to GRCm38 (mm10).

## Results

### Characteristics of congenic strains

We studied 1,293 congenic mice in total from 22 strains (Table 1, **S4 Table**) and confirmed the residual heterozygosity of these newly created strains was low (<0.2%; **S4 Figure**). The average adipose depot weight of the adult congenic mice ranged from 0.32±0.14 grams to 0.99±0.58 grams, depending on strain.

**Table 1.** Characteristics of congenic mice by strains

| Strain<sup>a</sup> | N | Age at necropsy (days) |  | Gonadal adipose (grams) | Body weight (grams) |
| --- | --- | --- | --- | --- | --- |
|  |  | Range | Mean±SD<sup>b</sup> | Mean±SD | Mean±SD |
| 1 | 113 | 175-187 | 180±2 | 0.54±0.24 | 31.77±2.93 |
| 1.1 | 176 | 162-184 | 180±2 | 0.55±0.28 | 31.44±2.96 |
| 1.1.1 | 3 | 180-180 | 180±0 | 0.99±0.58 | 36.13±4.72 |
| 1.2 | 14 | 180-182 | 181±1 | 0.61±0.28 | 33.43±4.24 |
| 3 | 175 | 177-190 | 180±2 | 0.56±0.24 | 31.18±3.19 |
| 3.1 | 120 | 178-182 | 180±1 | 0.55±0.25 | 31.64±3.21 |
| 3.1.1 | 106 | 178-186 | 180±1 | 0.50±0.23 | 31.55±3.11 |
| 3.1.1.1 | 127 | 179-184 | 181±1 | 0.39±0.17 | 31.10±2.26 |
| 3.1.1.2 | 34 | 179-181 | 180±1 | 0.40±0.17 | 31.21±2.58 |
| 3.1.1.3 | 14 | 179-181 | 180±1 | 0.56±0.38 | 31.23±3.77 |
| 3.1.1.4 | 63 | 103<sup>c</sup>-193 | 160±29 | 0.32±0.14 | 29.57±2.63 |
| 3.1.2 | 21 | 180-183 | 181±1 | 0.51±0.24 | 31.70±2.68 |
| 3.1.3 | 6 | 178-183 | 181±2 | 0.40±0.14 | 30.03±2.46 |
| 3.1.4 | 11 | 179-181 | 180±1 | 0.75±0.28 | 31.79±3.82 |
| 3.1.4.1 | 26 | 179-191 | 181±3 | 0.51±0.22 | 32.28±2.20 |
| 4 | 119 | 175-188 | 180±2 | 0.52±0.22 | 31.09±3.05 |
| 4.1 | 44 | 179-231 | 182±8 | 0.46±0.18 | 30.50±2.20 |
| 4.1a | 22 | 177-190 | 180±3 | 0.51±0.19 | 31.31±2.42 |
| 4.2 | 16 | 180-181 | 180±0 | 0.52±0.18 | 30.90±2.51 |
| 4.3 | 18 | 179-183 | 181±1 | 0.51±0.18 | 31.11±2.55 |
| 4.4 | 61 | 178-193 | 181±3 | 0.48±0.28 | 30.95±2.89 |
| 4.5 | 4 | 179-181 | 180±1 | 0.90±0.45 | 34.89±4.53 |

<sup>a</sup>Substrain IDs. <sup>b</sup>SD=0 indicates less than one day. <sup>c</sup>We necropsied 10 younger mice ranging from 103 to 124 days of age.

### Marker association analysis

In an initial general linear model analysis of a combined group of mice from all congenic strains, we estimated effect of marker genotype on gonadal adipose depot weight (Figure 2). The result confirmed a linkage with a large effect size at ~52.5 Mb; this effect size was 0.5 unit of pooled standard deviation for two independent genotype groups (Figure 2C), and 129-derived allele increase the phenotype (Figure 2B). However, the result also indicated possible additional peaks (Figure 2A).

**Figure 2.**
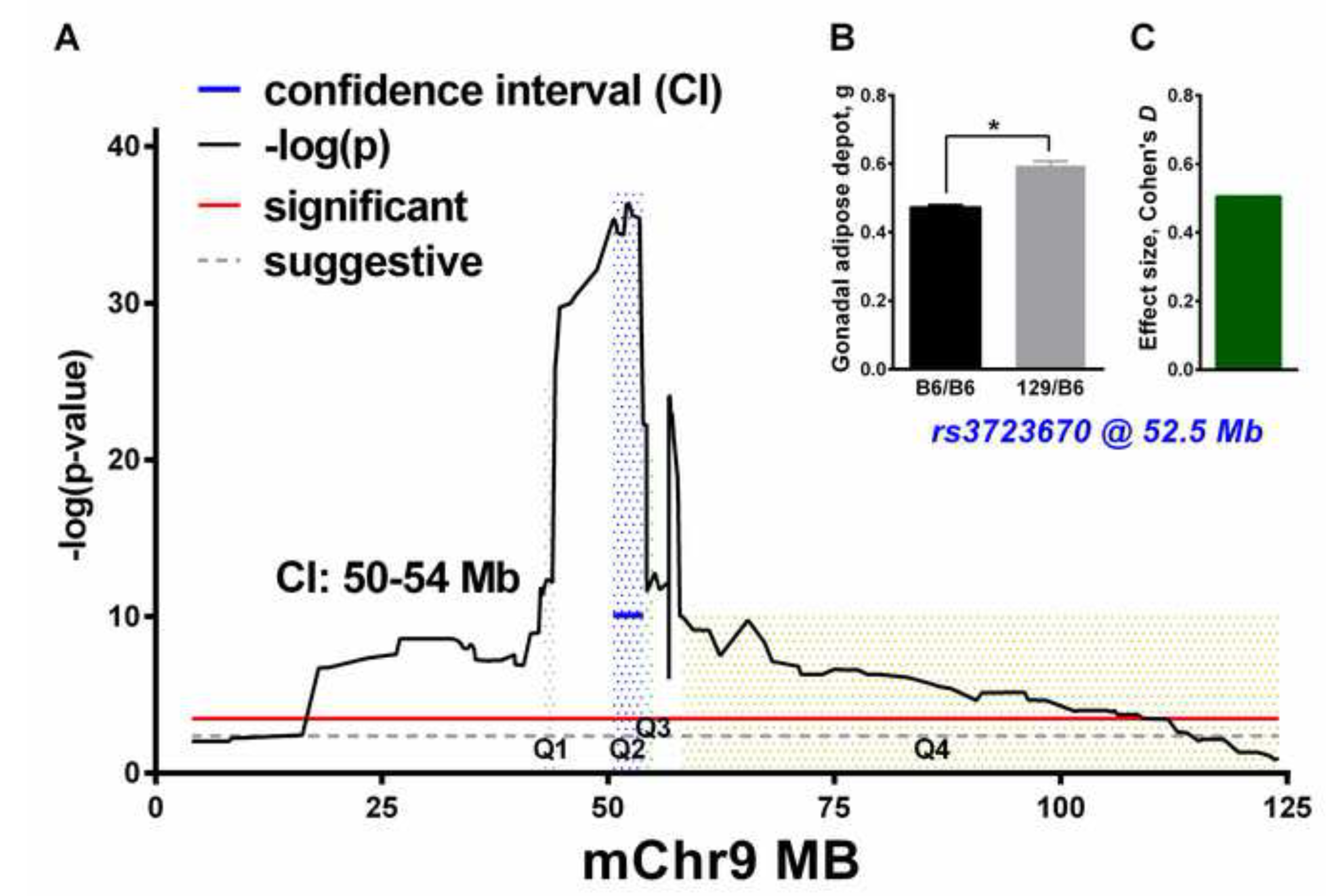
Detection of gonadal adipose depot weight QTLs on chromosome 9 by association analyses of a pooled congenic population. (**A**) Location of the markers in Mb on mouse chromosome 9 (mChr9); the y-axis is the -log p-values (black line) obtained in a general linear regression model analysis with body weight and strain as covariates (red line, significant threshold; gray line, suggestive). We defined the QTL confidence intervals by two units of -log10 p-value drop from the peak (blue line). For QTL1-QTL4 (Q1-Q4), the shaded peak areas correspond to QTL regions defined by the sequential method. (**B and C**) Average weight of the gonadal adipose depot by genotype (B6/B6, B6/129; **B**) and Cohen’s *D* effect size (**C**) of *rs3723670*, the most associated marker (i.e., the marker with the lowest p value). *p<0.0001, difference between genotypes.

### Common segment method

As expected, the common segment method was uninformative for both the narrower and broader analyses, failing to point to one region with a single QTL. In the narrow analysis, of the 12 congenic strains we evaluated, three were *Adip20(*+) and nine were *Adip20(*-) (**S5-6 Tables**). The positive strains had no single region in common. For the broad analysis, we used mice from 19 of the 22 strains (three strains had too few mice of a particular genotype for statistical analysis; **S4, S8 Table**); of these strains, five were *Adip20(*+) and fourteen were *Adip20(*-) (**S7-8 Tables)**. Like the narrow analysis, the broad analysis indicated that the positive strains had no single region in common.

### Sequential method

Using the sequential method, we analyzed the congenic strains with group sizes N ≥ 12/genotype/strain (**S5 Table**). We put the 12 congenic strains that met this sample-size criterion into one of three groups for comparison; the groupings and comparison order (described below) were suggested by the results of the MST analysis (Figure 3A). Within these groups, we compared pairs of strains, in the predetermined order, starting from comparison of the host strain (all mice without the donor fragment) and the strain with the smallest donor region (Figure 3B). The sequential analysis results indicated the presence of four QTLs (Figure 3C), which we ordered from proximal to distal, starting with QTL1. For most of the QTLs, the 129-derived allele increased the weight of the gonadal adipose depot, but QTL3 had the opposite direction of effect (Figures 3C and 4).

**Figure 3.**
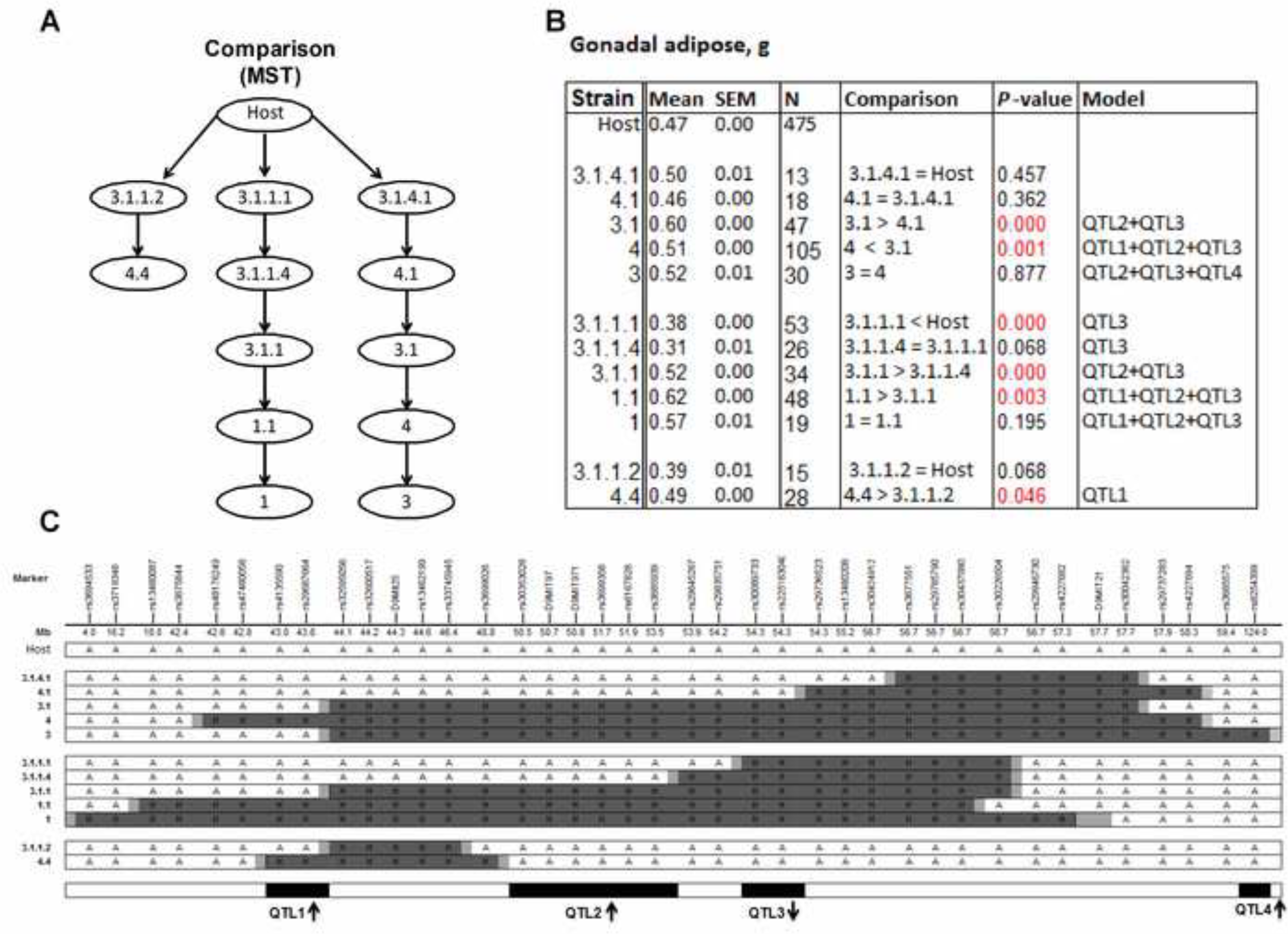
Dissection of QTLs with the congenic strains and the sequential method. (**A**) Minimum spanning tree (MST). (**B**) Mean and standard error of the mean (SEM) by strain for gonadal adipose depot weight in grams. Host = combined group of homozygous littermates without the donor fragment from all congenic strains; N = number of mice in each group. P-values are for least-significant-difference t-tests for each sequential comparison; red text indicates p-values that met the threshold (p<0.05). For the comparisons of two strains, the first strain was either greater (>), less than (<), or equal to (=) the second strain in gonadal adipose depot weight. “Model” lists QTLs within the donor fragment of each congenic strain suggested by the sequential analysis. (**C**) Congenic strains and genotypes for informative markers: A = B6/B6, shown as white bars; H = 129/B6, shown as dark gray bars; the recombinant ends correspond to both sides of the donor fragment for each congenic strain between H and A (or chromosome end), shown as light gray bars. QTLs are shown at the bottom as black boxes; arrows show the direction of effect (e.g., ↑ indicates that the QTL allele from the 129 strain increases the trait value). The data support the presence of four QTLs; we reported the detailed logic for this conclusion in **S1Text**.

**Figure 4.**
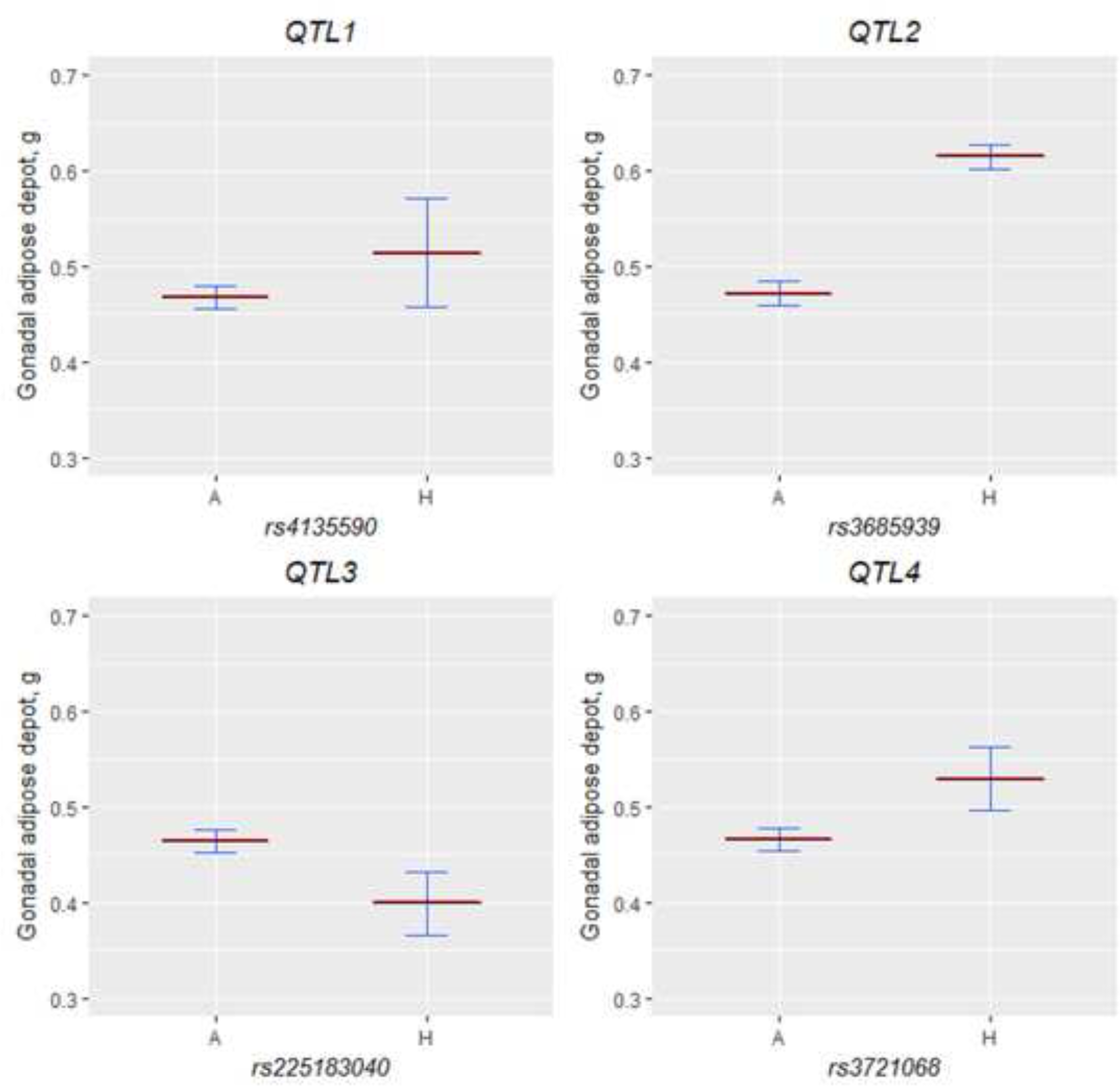
QTL effects. The least square means (dark red lines) and 95% lower and upper confidence limits (blue lines) for QTL genotypes were obtained from the general linear model with body weight as covariate in the congenic mouse data used for the sequential analyses. We selected one marker in the middle of each QTL to plot each QTL effect. The host (A) comprised littermates homozygous for B6/B6 from all strains; we selected appropriate congenic mice (H) as suggested by the results of the sequential analysis.

### Genetic variations

We analyzed genes residing in the QTL regions estimated in the sequential congenic analysis: QTL1, 9:42779420-44142555; QTL2, 9:48789608-53884418; QTL3, 9:54242579-54286770; and QTL4, 9:58262383-124595110. We refined the poorly localized QTL4, drawing on marginally significant mapping results from the original F_2_ intercross [15], to marker *rs3721068* with a 10-Mb flanking region (QTL4, 9:108915010-118915010). There were 44,009 variants between the 129P2/OlaHsd and C57BL/6J strains (closely related to the B6 and 129 parent strains used in our study), which we extracted (**S3 Figure**). Of these variants, the *in silico* analysis predicted that 2% change some aspect of mRNA regulation (**S9 Table**). There are 429 protein-coding genes [21], and 31% of the variants that map within the coding sequence are missense variants. Specifically, 48 of these genes contain either missenses or nonsenses variants (**S10 Table**).

### Previously reported mouse and human QTLs

Using these QTL co-ordinates, we surveyed the mouse genome database [4] for other adiposity QTLs on chromosome 9. Of the eight previously reported QTLs (*Adip5*, *Adip14*, *Dob2*, *Mobq8*, *Obq5*, *Carfhg2*, *Plbcq5*, and *Tdmq1*), seven QTLs had overlapping confidence intervals with the QTLs reported here (**S11 Table**). We also identified the regions of conserved synteny of human chromosomes 3, 11, and 15. We next examined results of recent human genome-wide association studies to find genes associated with body mass index and adiposity [28] and residing in these regions (Table 2), and to examine mouse orthologous genes with sequence variants. Four human genes (*FDX1*, *ARHGAP20*, *PDCD6IP,* and *TRAK)* associated with adiposity and six (*C11orf53*, *NCAM1*, *DMXL2*, *OSBPL10*, *LOC101928114,* and *CCK*) associated with body mass index (a commonly used but indirect measure of human adiposity) resided in these regions. Among mouse orthologues of these 10 genes, at least one, *Osbpl10*, has a protein-coding variant between 129P2/OlaHsd and C57BL/6J strains closely related to the B6 and 129 parental strains used in our study.

**Table 2.** Human genome-wide associations within mouse QTLs for adiposity and body mass index

| QTL | Mouse QTL location | Human region | Adiposity | Body mass index |
| --- | --- | --- | --- | --- |
| 1 | chr9:42779420-44142555 | chr11:119308720-120693788 | None | None |
| 2 | chr9:48789608-53884418 | chr11:107691025-114109156 | *FDX1, ARHGAP20* | *C11orf53, NCAM1* |
| 3 | chr9:54242579-54286770 | chr15:51295990-51341931 | None | *DMXL2* |
| 4 | chr9:108915010-118915010 | chr3:27712199-48621406 | *PDCD6IP, TRAK1* | *OSBPL10, LOC101928114, CCK* |

This table lists chromosome: bp locations for mouse (GRCm38; mm10) and human (GRCh38). We obtained human orthologous regions by the UCSC genome converter [90]. “Adiposity” and “Body mass index” columns contain Human Gene Nomenclature Committee gene symbols.

### Microarray results

The donor region of the congenic strain 4 (42.6 to 58.3 Mb; Figure 3C) has 277 genes, and in each microarray experiment, ~15% of these genes passed the microarray differential expression filters with a 1.5-fold change and/or a false discovery rate of p<0.05 (Figure 5). More than half of these genes were differentially expressed in both microarray Experiments 1 and 2 (*Abcg4, Acsbg1, Alg9, Arcn1, C230081A13Rik, Cd3g, Cryab, Dixdc1, Fam55d, Fxyd6, Hmbs, Il10ra, Isl2, Mcam, Mpzl2, Mpzl3, Nnmt, Ptpn9, Sik2, and Sik3;* Figure 5 and **S12 Table**). Two genes that were differentially expressed had two missense variants apiece (*Dixdc1* and *Sik2;* **S10 Table**). Five genes including two with the missense variants contained putative cis-regulator variants (*Alg9, Cryab, Dixdc1, Mcam*, and *Sik2*; **S9 Table**). We also confirmed the expectation that differentially expressed genes cluster in the congenic region rather than elsewhere in the genome (**S4 Figure**).

**Figure 5.**
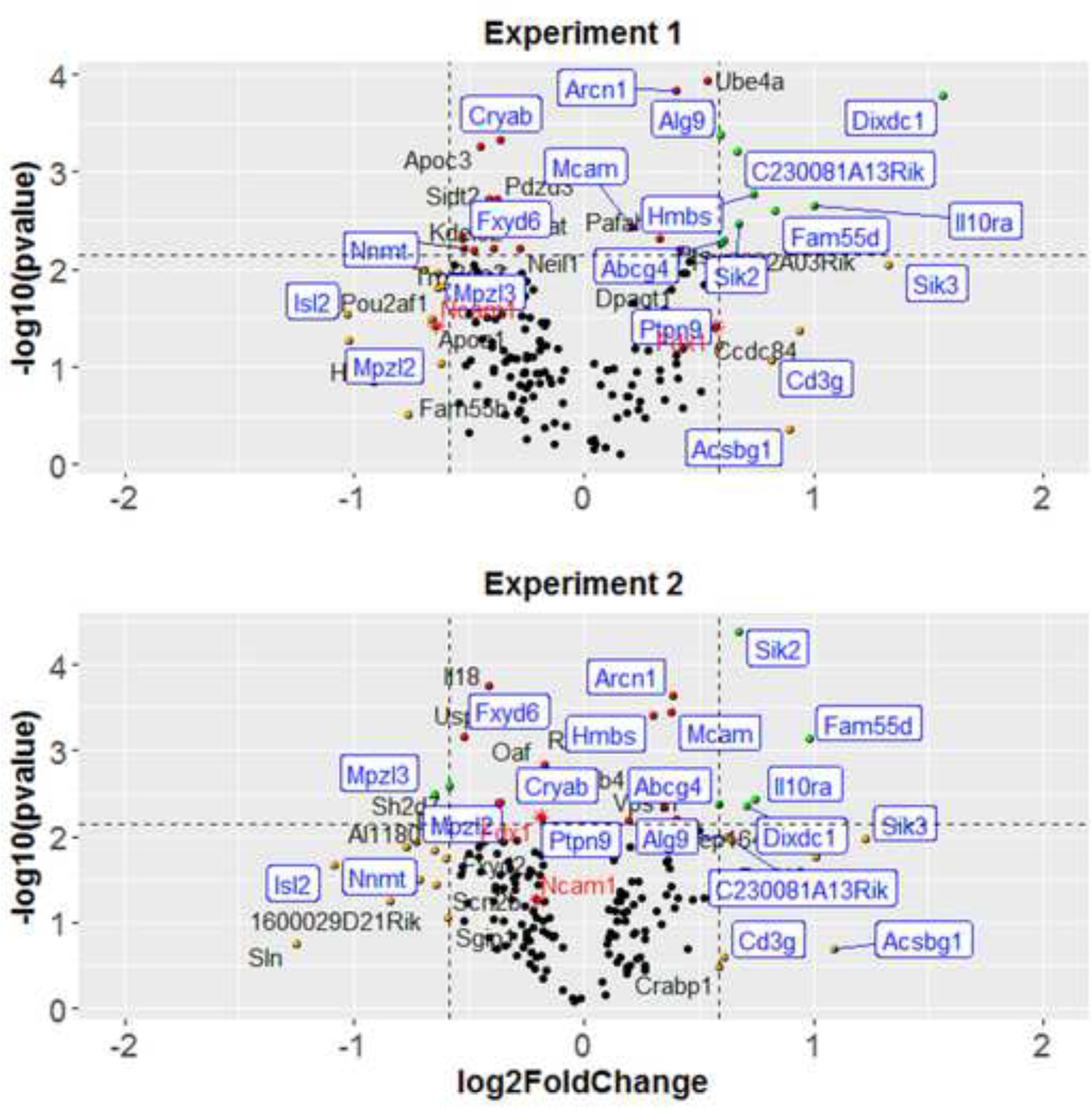
Extraction of differentially expressed genes: filtering on a volcano plot of the microarray data analysis of the Experiment 1 (top) and Experiment 2 (bottom). In Experiment 1, we selected the gonadal adipose tissue from male mice from congenic strain 4 with the donor region captured QTL1, QTL2, and QTL3 (42.6 to 58.3 Mb). Half of the mice were heterozygous for the donor region (129/B6; N=7), and half were littermates homozygous for the donor region (B6/B6; N=7). Experiment 2 replicated Experiment 1 except with N=6 mice in each genotype group, for a total of 12 mice. The customized volcano plots depict the estimated log_2_-fold change (x-axis) and statistical significance (−log_10_ p-value; y-axis). Each dot represents one gene. The green dots show genes with absolute log_2_-fold change > 0.58 (i.e., 1.5-fold change) and false discovery rate < 0.05; the red dots show genes with false discovery rate < 0.05; the yellow dots show genes with absolute log_2_-fold change > 0.58; the black dots show genes that did not reach these statistical thresholds. We label genes that passed any filters by name. Twenty differentially expressed genes are reproducible between Experiment 1 and 2, labeled with blue text and square boxes. Red stars show two genes that passed the filters in one but not both experiments yet are noteworthy because their human orthologues are associated with adiposity or body mass index in genome-wide association studies (Table 2). We summarize the additional experimental details in **S12 Table**.

Integrating all methods above, there was no convergence to a single candidate gene (Figure 6). Several genes did not survive the candidate gene filters used here but are noteworthy near misses because their human orthologues are associated with similar traits in genome-wide association studies (*Fdx1* and *Ncam1*; Table 2). While no single gene was implicated consistently, several had more evidence than did others, for instance, *Dixdc1* and *Sik2*, as mentioned above, were differentially expressed in gonadal adipose depots and contained several putative regulatory and missense variants.

**Figure 6.**
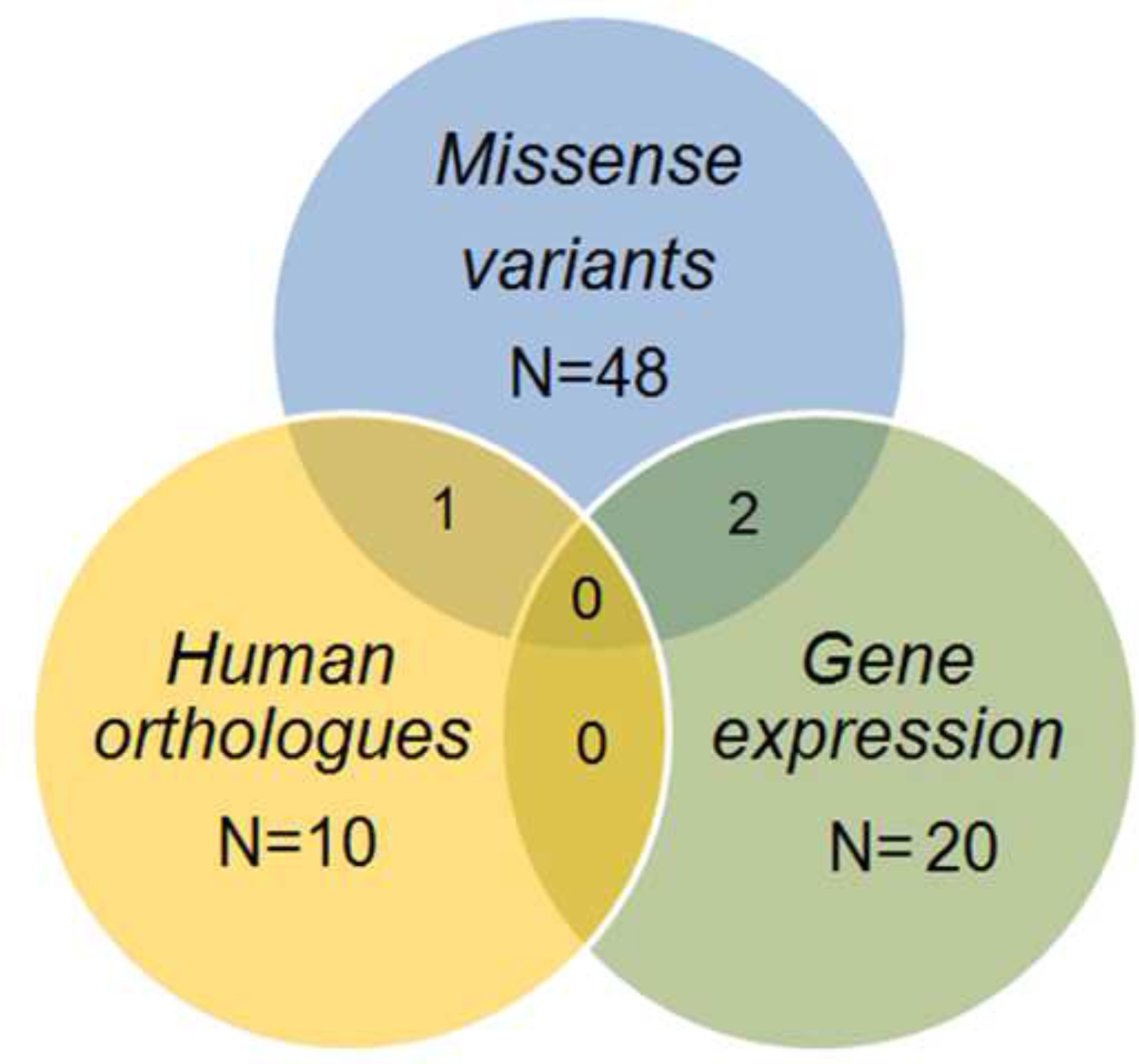
**The overlap of** candidate gene by nomination method. We extracted candidate genes based on evidence from missense genetic variation, human adiposity, or body mass index orthologues and differential gene expression.

## Discussion

Here we bred congenic strains with the expectation that we could narrow the genomic location of an adiposity QTL, *Adip20*, to a region with only one or a very few genes on mouse chromosome 9. The successful identification of several obesity genes, including leptin [39], the leptin receptor [40], tubby [41, 42], and agouti [43], suggested this strategy would be useful even when effect sizes differ by genetic context [44]. These early successes spawned other efforts like ours to map genetic variants that contribute to obesity [45–62], and investigators have identified hundreds of QTLs [4]. When we fine-mapped the *Adip20* locus, we detected effects of at least four QTLs on chromosome 9. These results illustrate both the complexity of body composition genetics and the utility of congenic strains to better characterize the underlying genetic architecture and, further, suggest that decomposition of a single QTL into multiple QTLs is commonplace for traits like adipose depot weight and obesity [63–65].

In general, the study of complex traits is easier when we reduce or eliminate extraneous sources of trait variation. For obesity, sex-by-genotype interactions are a common feature of natural populations, including humans [66], as well as mouse intercross studies [8, 67, 68]. Thus, we chose to study only male mice to reduce the potential interaction of a particular genotype with sex, while acknowledging this choice makes the results less generalizable [69]. In addition to reducing variation in adiposity due to sex, we controlled age as a source of potential trait variation to reduce genotype-by-age interactions [70]. We also tried to reduce parent-of-origin and maternal effects because of their known influence on adiposity [71, 72]. We controlled some maternal effects in the sense that all congenic mice had mothers of the same genotype, and we controlled some imprinting and other parent-of-origin effects because all congenic mice inherited the donor region from the father (i.e., paternal transmission). We acknowledge that maternal effects are a composite of many factors, not just maternal genotype, and so the offspring may vary based on maternal body weight and parenting behavior. However, generally, these experimental choices were useful in reducing certain types of variation.

Obesity as a trait is both simple and complex: it is simple conceptually because its single hallmark is an excess of adipose tissue. However, it is complex because not all adipose tissue is the same. It is stored in depots that differ in location, size, and function [13, 14, 73], and each depot has a different genetic architecture [15] and pattern of gene expression [74, 75]. Further complexities arise because adipose tissue is hard to measure directly, and each measurement method yields different results [76]. Here we chose to study one adipose depot, which we measured directly by necropsy. This is a simple measurement method, and our expectation was that its genetic architecture would be less complex than for whole-body obesity. We specifically chose the gonadal over other depots because it is the heaviest [13, 14, 73] and a proxy for overall adiposity [77, 78]. However, despite all these complexities, both mouse and human genetic approaches point to the same underlying mammalian biology: QTL regions found here contain many mouse orthologues for obesity genes identified by human genome-wide association studies.

We began the construction of congenic mice guided by mapping results from the intercross population [15]. We also bred reciprocal consomics, which helped us define the genetic architecture [11], and provided a source of breeder mice for constructing the congenic strains. Building on information from the consomic mice study [11], we learned that a single *Adip20* QTL on chromosome 9 could not explain the observed results from the reciprocal strains [11]. Study of the congenic mice here confirmed and extended those more general results, uncovering the location of the multiple QTL and identifying their opposing effects.

Within the four QTL regions we identified, there are many candidate genes, and we see a technical gap between available and ideal testing methods. The most common available method is the total elimination of each gene (e.g., gene knockout) and quantification of knockout versus control mice. However, this method has a serious limitation as an approach to study obesity: large-scale knockout surveys indicate that inactivating mutations affect early embryonic growth for nearly half of genes tested [79], and about a third of genes affect body composition in adult mice [80, 81]. This method has poor specificity, so we cannot assume that if knockouts differ from controls in body composition (e.g., [82]), then the target gene is causal. Likewise, bioinformatics methods to evaluate functional variants are also not ideal because so many genes have potentially functional variants. Furthermore, we cannot completely exclude any of the thousands of observed variants because evaluating potential function with currently available informatics methods remains imperfect. For complex traits, the challenge is now to find new ways to winnow a large number of variants to only a few or to find a way to test many variants and their combinations rapidly. While there are statistical methods to winnow the candidate gene by trying to parse causal relationships among genotype gene expression and trait [83–85], ultimately we need direct tests of individual genotypes on traits.

While congenic mapping methods were useful in identifying additional adiposity QTLs on mouse chromosome 9, there are at least three limitations to consider when interpreting the results. The study of male mice exclusively we mentioned earlier. A second limitation is the residual heterozygosity within the congenic strains that we confirmed through genotyping. The amount was low relative to the number of backcross cycles because we used marker-assisted breeding, but still it may have contributed to experimental noise. To put this limitation in a broader context, we have learned recently from whole-genome sequencing studies that this approach will not ensure that all members of a particular inbred strain are completely genetically identical [86]. Whether variants among members of the same inbred strain are technical artifacts or represent true diversity is controversial, but if true this may explain in part the rapidly divergence in body weight during selective breeding of inbred mouse strains [87]. The third limitation is that the congenic breakpoints were concentrated more in the middle regions of the chromosome, and we may have missed additional QTLs in the proximal region where there were fewer breakpoints.

We used complementary methods to evaluate causal genes, but no single gene emerged as a highly prioritized candidate within any QTL region. This lack of convergence among methods may be due in part to the differing assumptions of each. For instance, the microarray approach we used here assumes that causal variation affects gene expression in the gonadal adipose depot of adult mice, but the actual causal variants may not affect gene expression at all or may act in a different tissue or at a different time in development. Obviously, the limitation of microarray experiments is the target tissue selection, and we acknowledge that genotype effects in other tissue could be causal (e.g., liver tissue [84, 88]). These points made, there was some convergence among methods. Specifically, the *OSBPL10* gene was identified by a human genome-wide association study of body mass index [89], and the mouse orthologue of this gene, which is within the QTL4 region, had a missense variant between the 129 and B6 mouse strains. Similarly, mouse gonadal adipose tissue expresses different amounts of *Dixdc1* and *Sik2* mRNA, depending on QTL2 genotype, and each of the corresponding genes has two predicted missense and several regulatory variants. These results provide direction in the future evaluation of candidate genes.

## Acknowledgements

We gratefully acknowledge the assistance with animal breeding of Rebecca James, Liang-Dar (Daniel) Hwang, Zakiyyah Smith, Matt Kirkey, Amy Colihan, and Laurie Pippett. We also acknowledge Richard Copeland and the consistent high-quality assistance of the animal care staff at the Monell Chemical Senses Center and thank them for their service. Michael G. Tordoff and Gary K. Beauchamp commented on a draft of the manuscript. Joseph H. Nadeau provided valuable advice on analytical approaches and manuscript preparation. We thank the two anonymous reviewers for their constructive comments on this manuscript.

## Funding

National Institutes of Health grants R01 DK094759, R01 DK058797, P30 DC011735, S10 OD018125, S10 RR025607, S10 RR026752, and G20 OD020296 and institutional funds from the Monell Chemical Senses Center funded this work, including genotyping and the purchase of equipment. The Center for Inherited Disease Research though the auspices of the National Institutes of Health provided genotyping services.

## Accession ID

We used mice for these experiments, NCBI Taxon ID 10090.

## Electronic Resources

http://www.cidr.jhmi.edu/nih/Completed_Projects_Report_June15.pdf

## Supporting Information

**S1 Table.** Markers used for genotyping and their physical positions on chromosome 9

**S2 Table.** List of mice with tumors or otherwise excluded from analysis Weight = Body weight in grams; Length, = body length in centimeters; Gonadal = gonadal adipose depot weight at the time of dissection in grams; Tumor location includes, weights if available

**S3 Table.** Statistics for descriptive and normality tests of gonadal adipose depot weight in congenic populations

**S4 Table.** Weight of gonadal depot and body weight (BW) of congenic mice with donor regions of different genotypes of donor regions ^a^Donor region genotype: Host = B6/B6; Donor = 129/B6. ^b^Donor region size: NA = mice without donor region; Partial = mice that inherited a partial-length (due to a recombination event) donor region from father; Full = mice that inherited full-length donor region from the father

**S5 Table.** Genotypes and phenotypes of the 12 congenic strains (strains with numbers of mice ≥12 mice/strain, and all mice having full-length donor regions) Column headings show IDs of congenic strains. We show here only markers informative for breakpoints of the donor regions. A=B6/B6 genotype; H=129/B6 genotype; X=data are not available. Based on analysis of gonadal depot weight (**S4,6 Table**), we identified strains as Adip20(+) (shaded; strains with significant differences between mice with B6/B6 and 129/B6 genotypes) or Adip20(-) (no shading; strains with no significant differences between mice with B6/B6 and 129/B6 genotypes). The far right column ("Shared region") indicates whether a QTL-containing region is shared only by Adip20(+) but not by Adip20(-) strains (i.e., whether a marker genotype is H in all Adip20(+) strains and A in all Adip20(-) strains). None of the regions of chromosome 9 met this single QTL only criterion.

**S6 Table.** Narrow analysis of gonadal depot weight in the 12 congenic strains (with numbers of mice ≥12 mice/strain, and all mice having full-length donor regions): general linear model (GLM), least square means, and post hoc Fisher's least significance difference (LSD) tests. N=Numbers of mice in each group with and without a donor region (donor genotype 129/B6; host genotype B6/B6). The donor group includes only mice with full-length length donor regions.

**S7 Table.** Broad strain inclusion criterion. Genotypes and phenotypes of the congenic strains of any sample size, except as noted below, including mice with partial or full-length donor regions. However, we still excluded three strains from this analysis (shaded in light grey) because they had very few mice (1.1.1, 3.1.3, 4.5). For other details, see **S5 Table**.

**S8 Table.** Broad analysis of gonadal depot weight using the criterion from **S7 Table**.

**S9 Table.** Regulatory variants within the four QTL regions predicted with an on-line tool Variant Effect Prediction[25]

**S10 Table.** Sorting Intolerant From Tolerant (SIFT) prediction of missense and stop codon gain/loss sequence variants of genes residing within the 4 adiposity QTL regions on chromosome 9

**S11 Table.** Summary of the mouse adiposity QTLs on chromosome 9 Type = mapping population; Con = congenic; Overlap refers to QTL1-QTL4 as identified by the sequential method. Approach = method of estimating confidence interval (LDR = logarithmic transformed probability; LOD = logarithm of the odds); Ref=reference.

**S12 Table.** Filtering of microarray data to identify reproducible candidate genes ^a^There are 277 genes within the donor region of strain 4 (42.6 to 58.3 Mb). Genes that passed the filters for differential expression in both Experiment 1 and 2 we display with blue font. Two genes in red font miss this criterion but their human orthologues are associated with adiposity or body mass index in GWAS studies (see **Table 2**). We show the remaining genes in black font.

**S1 Text**. Logic of the sequential method for each of the three branches of the minimum spanning tree (MST; central, left, and right) in Figure 3. Arrows after QTLs indicate the direction of effect of the 129-derived allele.

**S1 Figure.** Cullen and Frey graphs and empirical and theoretical data for untransformed and log-transformed congenics data. We tested the distribution of the gonadal adipose depot weight data for normality; **S3 Table** contains the parameters of the model-fitting untransformed and log-transformed data. CDF = cumulative distribution function.

**S2 Figure.** Pearson correlations between (A) gonadal adipose depot and body weight, (B) gonadal adipose depot and age as well as (C) body weight and age of all congenic mice. Gonadal = Gonadal adipose depot in grams; BW = body weight in grams; Age in days.

**S3 Figure.** Predicted effect for variants within four QTLs. In total, we uploaded 44,009 variants for the variant function analyses to the online tool Variant Effect Predictor; we found data from nearly all variants (99.8%) using this tool; of these variants, 2% were regulatory variants, and 31% of variants in coding regions were missense variants.

**S4 Figure.** Genome-wide overview of the microarray data from Experiment 1 with differentially expressed genes (blue dots) in adipose tissue between genotypes (129/B6 vs B6/B6) in congenic male mice. We observed potential heterozygosity (green dots) of this congenic strain in at least one of the samples at 25 of the 2,715 polymorphic markers. Of these, 19 were located in the congenic donor region (red dots), leaving 7 markers (green dots) with potential residual heterozygosity that may have produced differential gene expression independent of the donor region.

Click here to access/download

Supporting Information

[Supplemental Figure 1.docx](http://www.editorialmanager.com/pone/download.aspx?id=21150386&guid=ce825d32-4028-48ab-b1b4-a8efd3e71a88&scheme=1)

Click here to access/download

Supporting Information

[Supplemental Figure 3.docx](http://www.editorialmanager.com/pone/download.aspx?id=21150388&guid=d0ca9d74-7a8b-4799-96b8-3c6ada2ac954&scheme=1)

Click here to access/download

Supporting Information

[Supplemental Figure 4.docx](http://www.editorialmanager.com/pone/download.aspx?id=21150389&guid=1af0d1b4-f9e6-46a0-968c-f9961278f35e&scheme=1)

Click here to access/download

Supporting Information

[Supplemental Table 1.xlsx](http://www.editorialmanager.com/pone/download.aspx?id=21150390&guid=c254193b-d6a7-4045-932e-48450cf4a860&scheme=1)

Click here to access/download

Supporting Information

[Supplemental Table 2.xlsx](http://www.editorialmanager.com/pone/download.aspx?id=21150391&guid=a1a8b08d-035c-426b-b91f-f5ec2dce4a36&scheme=1)

Click here to access/download

Supporting Information

[Supplemental Table 3.xlsx](http://www.editorialmanager.com/pone/download.aspx?id=21150392&guid=ea7e1c4e-03a7-4819-b900-dc59eff3e3dc&scheme=1)

Click here to access/download

Supporting Information

[Supplemental Table 4.docx](http://www.editorialmanager.com/pone/download.aspx?id=21150393&guid=8c4d4d0e-c9b6-498a-9606-b3d5e1f19d39&scheme=1)

Click here to access/download

Supporting Information

[Supplemental Table 5.docx](http://www.editorialmanager.com/pone/download.aspx?id=21150394&guid=66c34dcf-7b10-4c58-ad76-f24a461bca74&scheme=1)

Click here to access/download

Supporting Information

[Supplemental Table 6.xlsx](http://www.editorialmanager.com/pone/download.aspx?id=21150395&guid=880cacba-f24f-4385-9b9b-f1c079d2b4bc&scheme=1)

Click here to access/download

Supporting Information

[Supplemental Table 7.docx](http://www.editorialmanager.com/pone/download.aspx?id=21150396&guid=cc1fcc6a-4d49-4a87-9e69-8e4a1af789e4&scheme=1)

Click here to access/download

Supporting Information

[Supplemental Table 8.xlsx](http://www.editorialmanager.com/pone/download.aspx?id=21150397&guid=9cacf5a6-8274-47fe-92ab-4e1350e6a71b&scheme=1)

Click here to access/download

Supporting Information

[Supplemental Table 9.xlsx](http://www.editorialmanager.com/pone/download.aspx?id=21150398&guid=fe542de1-314c-4ee5-8b5e-6de307176dd6&scheme=1)

Click here to access/download

Supporting Information

[Supplemental Table 10.xlsx](http://www.editorialmanager.com/pone/download.aspx?id=21150399&guid=dffaa9da-28ab-4185-8a53-105b77618bcd&scheme=1)

Click here to access/download

Supporting Information

[Supplemental Table 11.docx](http://www.editorialmanager.com/pone/download.aspx?id=21150400&guid=fae808e0-9c2a-42d5-a462-f468f2acdb65&scheme=1)

Click here to access/download

Supporting Information

[Supplemental Table 12.docx](http://www.editorialmanager.com/pone/download.aspx?id=21157151&guid=878f70e8-e909-498f-a912-c0b37df90a1e&scheme=1)

Click here to access/download

Supporting Information

[Supplemental Text 1.doc](http://www.editorialmanager.com/pone/download.aspx?id=21150402&guid=8e6051f6-6981-4f5e-817e-b7d08557fad6&scheme=1)

